# Tumor subclonal progression model for cancer hallmark acquisition

**DOI:** 10.1101/149252

**Authors:** Yusuke Matsui, Satoru Miyano, Teppei Shimamura

## Abstract

Recent advances in the methods for reconstruction of cancer evolutionary trajectories opened up the prospects of deciphering the subclonal populations and their evolutionary architectures within cancer ecosystems. An important challenge of the cancer evolution studies is how to connect genetic aberrations in subclones to a clinically interpretable and actionable target in the subclones for individual patients. In this study, our aim is to develop a novel method for constructing a model of tumor subclonal progression in terms of cancer hallmark acquisition using multiregional sequencing data. We prepare a subclonal evolutionary tree inferred from variant allele frequencies and estimate pathway alteration probabilities from large-scale cohort genomic data. We then construct an evolutionary tree of pathway alterations that takes into account selectivity of pathway alterations via selectivity score. We show the effectiveness of our method on a dataset of clear cell renal cell carcinomas.

## 1 Scientific Background

Cancer is a heterogeneous genetic disease characterized by dynamic evolution through acquisition of genomic aberrations. The clonal theory of cancer proposed by Nowell [1] postulates that acquisition of a mutation in cancer follows natural selection of the Darwinian model, in which cancer obtains the advantages of biological fitness under selective pressure.

The development of multiregional sequencing techniques has provided new perspectives on genetic heterogeneity [2]. According to studies on multiregional sequencing, spatially distinct regions within the same tumor acquire different sets of somatic single-nucleotide variants (SSNVs), and this phenomenon is called *intratumor heterogeneity*. Recently, methods for reconstruction of cancer evolutionary structures were extensively studied. Because the cell population of each region is a mixture of normal and tumor cells, distinct regions are deconvoluted into cell subpopulations called subclones, and then they are assigned to tree structures under the constraint that is derived from the infinite site assumption [3].

Nonetheless, identification of clinically actionable subclone targets for individual patients remains problematic. One of the reasons is that resulting subclonal trees are too diverse to interpret with clinical information. In this direction, Matsui *et al.* [4] proposed clustering methods for cancer evolutionary trees based on the tree shapes and interpreting clinical impact of subclonal evolution via clustering results with clinical information of each tree. Another reason is the difficulties with identifying the most plausible biological event from the limited number of SSNVs because of the sequencing depth and low frequencies of mutations.

With the aim to overcome this problem, we developed a novel method for inferring a tumor progression model in terms of acquisition of cancer hallmarks (Figure 1). Our contributions are as follows: (1) proposing a novel framework for interpreting the cancer evolutionary tree using pathways and cancer hallmarks (2) developing the integrative approach to estimate individual each subclone’s cancer hallmarks by merging multiregional sequencing data and large scale genomic cohort data. We also demonstrate the effectiveness of our method on an actual dataset of clear cell renal cell carcinomas (ccRCCs).

**Fig. 1.**
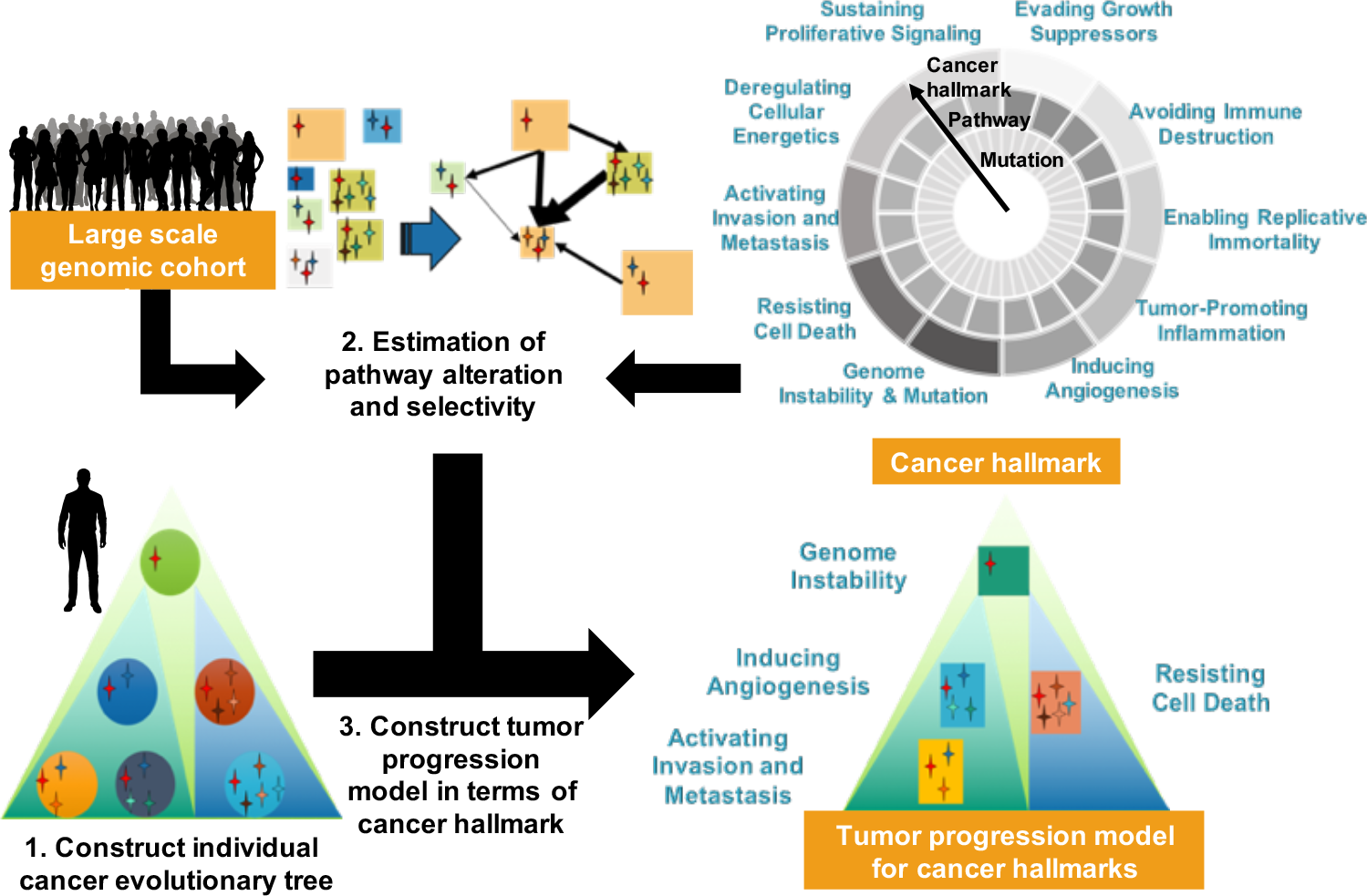
Overview of our proposed approach for inferring tumor progression model for cancer hallmarks to interpret the biological functions of each subclones. The proposed method consists of 3 steps: (1) constructing a cancer subclonal evolutionary trees for individuals by multiregional sequencing data, (2) estimating pathway alteration probabilities and the strength of selective pressure between pathway alterations, and (3) constructing a model of progression of pathway alterations with cancer hallmarks.

## 2 Materials and Methods

Our method consists of 3 steps: (1) constructing a skeleton of a cancer subclonal evolutionary tree, (2) estimation of pathway alteration probability and calculation of *selectivity score*, and (3) constructing a model of progression of pathway alterations.

First, we construct an a priori evolutionary tree of the pathway alteration progression model, called a skeleton, to decompose the cell population into subclones and infer the subclonal evolutionary structures for each patient on the basis of multiregional variant allele frequencies (VAFs). Second, we estimate the pathway alteration probability by means of a large cohort dataset to identify the most likely pathway alterations in the subclones and to calculate *selectivity score, i.e.*, the strength of the selectivity among the pathway alterations at the subsequent step. At the last step, we construct the tumor progression model of pathway alterations based on the skeleton and pathway alteration probabilities. After scanning each subclone to test whether at least 1 SSNV is included in the given pathways, we remove the subclones in the first case or we identify the unique pathway alteration from those pathways in the second case under the assumptions because there are cases of no candidate pathway alteration or multiple candidate pathway alterations per subclone.

**(Assumption 1)** No pathway alteration occurs twice in the course of cancer evolution.

**(Assumption 2)** No pathway alteration is ever lost.

Assumptions 1 and 2 mean that if a given pathway alteration occurs in any subclone, it happens exactly once in the course of tumor progression. We reconstruct the progression model from ancestral subclones, and we never use the pathway alterations in those subclones for their descendant subclones. In addition to the 2 assumptions, we assume selective pressures between the pathway alterations.

**(Assumption 3)** There is selective pressure between the pathway alterations.

We model this system on the basis of the notion of conditional probability, which can be estimated from large-scale cohort datasets. Using the 3 assumptions, we identify a unique pathway alteration from the multiple candidates of pathway alterations with the strongest selectivity. In the following section, we describe our method in more detail.

### 2.1 Constructing a skeleton of the cancer subclonal evolutionary tree

This skeleton represents the subclonal evolutionary tree based on VAFs obtained by bulk sequencing from a single patient with multiple regions. The VAFs are approximately proportional to the sizes of cell populations with the set of SSNVs; however, in the settings of bulk data, each region may be a mixture of normal and tumor cells and may require deconvoluting the cell populations into subpopulations. The identified subclones are assigned to tree structures via 2 assumptions: (i) a mutation cannot recur in the course of cancer evolution, and (ii) no mutation can be lost [6]. Several approaches are implemented to deal with the problem. Using one of the algorithms, LICHeE [3], we construct the skeleton from the VAFs for each patient. Let 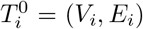 be a skeleton of patient *i*; *i* = 1, 2,…, *n* with a set of vertices *V*_*i*_ = {*ν*_*ij*_; *i* = 1, 2,…, *n*, *j* = 1, 2,…, *η*_*i*_} and edges *E*_*i*_ = {*e*_*ik*_; *i* = 1, 2,…, *p*, *k* = 1, 2,… *v*_*i*_}, where vertices and edges represent subclones with a set of SSNVs and evolutionary relations, respectively. Without a loss of generality, *ν*_*i*,*j*=1_ always represents normal cells. Each vertex has a set of labels that can be obtained from the mapping *L*: *ν*_*i*,*j*_ ↦ *L*_*i*,*j*_, e.g., *L*_*i*,*j*_ = {SSNV1, SSNV2, SSNV3}.

### 2.2 Estimation of pathway alteration probability and selectivity score

To carry out phenotypic characterization of each subclone, we need to identify the most closely related pathway alterations for subclones. There are mainly 2 approaches to detection of pathway alterations: one is knowledge based gene enrichment analysis such as Fisher’s exact test, and the other is a *de novo*-oriented approach, where the alteration patterns are mapped to large-scale protein networks and identify subnetworks as a driver pathway with cost functions such as a tendency for mutual exclusivity. In this study, we focus on the knowledge based approach because the biological validation for *de novo* pathways is usually difficult to perform quickly.

Using the large-scale cohort data, because of the nature of the limitation on sample size in experimental data, we estimate the pathway alteration probability using SLAPenrich [5], which is a state-of-the-art method for identifying pathway alteration and provides background pathway alteration probabilities and mutual exclusivity. Let *P* denote the pathway list, and then the main output of SLAPenrich is the *P*-value *p*_*i*_; *i* = 1, 2,…, |*P*| for each pathway *i* that represents the significance of the enrichment of mutations. Optionally, we can obtain pathway alteration probabilities for each sample, that is, *ρ*_*i*,*j*_ = Pr(*X*_*i*,*j*_ ≥ 1); *i* = 1, 2,…, |*P*|, *j* = 1, 2,…, *N* where *X*_*i*,*j*_ represents the number of SSNVs that are included in pathway *i* in sample *j*.

We define binary variable *y*_*i*,*j*_ if *ρ*_*i*,*j*_ > *t* then *y*_*i*,*j*_ = 1 otherwise *y*_*i*,*j*_ = 0 with a given threshold 0 ≤ *t* ≤ 1. We evaluate the selectivity score, *S*_*k*→*l*_; *k*, *l* = 1, 2,…, |*P*|, *k* ≠ *l*, defined as

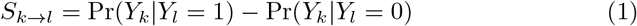

where *Y*_*k*_ represents a variable denoting the alteration status of pathway *k*. We empirically estimate *S*_*k*→*l*_ as follows:

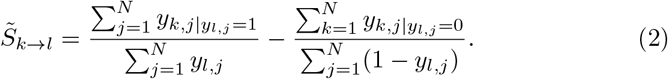

We consider zero or negative values to represent a no selectivity, i.e., 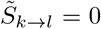 if 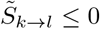.

### 2.3 Constructing a model of progression of pathway alterations

Now we are ready to construct the subclonal evolutionary tree for pathway alterations. Given a subclone, we first scan the SSNVs to identify candidate pathway alterations. If at least 1 SSNV is included in a pathway, this situation is called a *candidate pathway alteration*. Suppose *P*_*k*_; *k* = 1, 2,…, |*p*| is the genes included in pathway *k* and *Z*_*i*,*j*,*k*_ is the candidate pathway alteration status of subclone *j* in patient *i* where *Z*_*i*,*j*,*k*_ = 1 if *L*(*ν*_*i*,*j*_) ⊆ *P*_*k*_; *i* = 1, 2,…,*n*,*j* = 1, 2,…, *η*_*i*_ otherwise *Z*_*i*,*j*,*k*_ = 0. In case of *Z*_*i*,*j*,*k*_ = 0 for all *k*, we regard the subclone as a nonfunctional one and remove node *ν*_*i*,*j*_ and the corresponding edges from skeleton 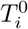.

By means of candidate pathway alterations *Z*_*i*,*j*,*k*_, we identify the unique pathway alteration. In the event that an ancestral subclone consists of normal cells, we select a pathway with the smallest *P*-value, i.e.,

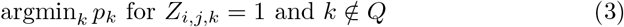

Otherwise, we select the pathway with the highest selectivity score, i.e.,

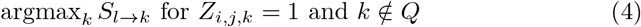

where *l* is the pathway alteration of the ancestral subclone, and *Q* is a set of pathway alterations that have already appeared in the ancestral subclones. Assumptions (1) and (2) are ensured by the condition *k* ∉ *Q*. If there is no corresponding pathway alteration because all the candidate pathway alterations have already taken place in the ancestral subclones, then we remove the subclone as a nonfunctional one.

### 2.4 The dataset

A dataset from a study on ccRCCs [7] was used for the analysis. Whole-exome multiregional bulk sequencing was performed on 8 individuals with clinical information, and 587 out of 602 mutations remained after filtering of mutations with depth less than 100 ×.

The estimation of pathway alteration probabilities followed SLAPenrich procedures described in [5]. The 417 KIRC (corresponding to ccRCC) samples from The Cancer Genome Atlas (TCGA) and International Cancer Genome Consortium (ICGC) and high-confidence variants identified in another study, [8], were used for estimation of the pathway alteration probabilities. Pathway gene sets were downloaded from the Pathway Commons data portal (v8, 2016/04), and gene sets containing fewer than 4 or more than 1,000 genes were discarded. After merging the gene sets that correspond to the same pathway across multiple data sources or have a large overlap defined as Jaccard index ≥ 0.8, we obtained 1,911 pathway gene sets. Cancer hallmarks were manually curated and assigned to 456 pathways [5].

## 3 Results

We reconstructed the skeletons from VAFs by means of LICHeE using the same parameters in their experimental settings that are described in [3], and eventually, 8 skeletons were obtained. Next, we estimated the pathway alteration probabilities based on SLAPenrich and obtained *P*-values for 209 pathways and pathway alteration probabilities for 417 patients. We evaluated the selectivity score with the threshold *t* = 0.1.

We show the results of the constructed models of ccRCC progression in terms of cancer hallmark acquisition in Figure 2. The complexity of cancer evolutionary trees were reduced because several SSNVs in subclones plays the similar roles in terms of the biological pathway. Specifically, EV003 were reduced to only the two pathways alteration, which means most of SSNVs in the subclones are involved in the TP53 related pathway. In this way, our approach could summarize the original cancer evolutionary trees and give biological interpretation via mapping SSNVs to the biological pathways.

**Fig. 2.**
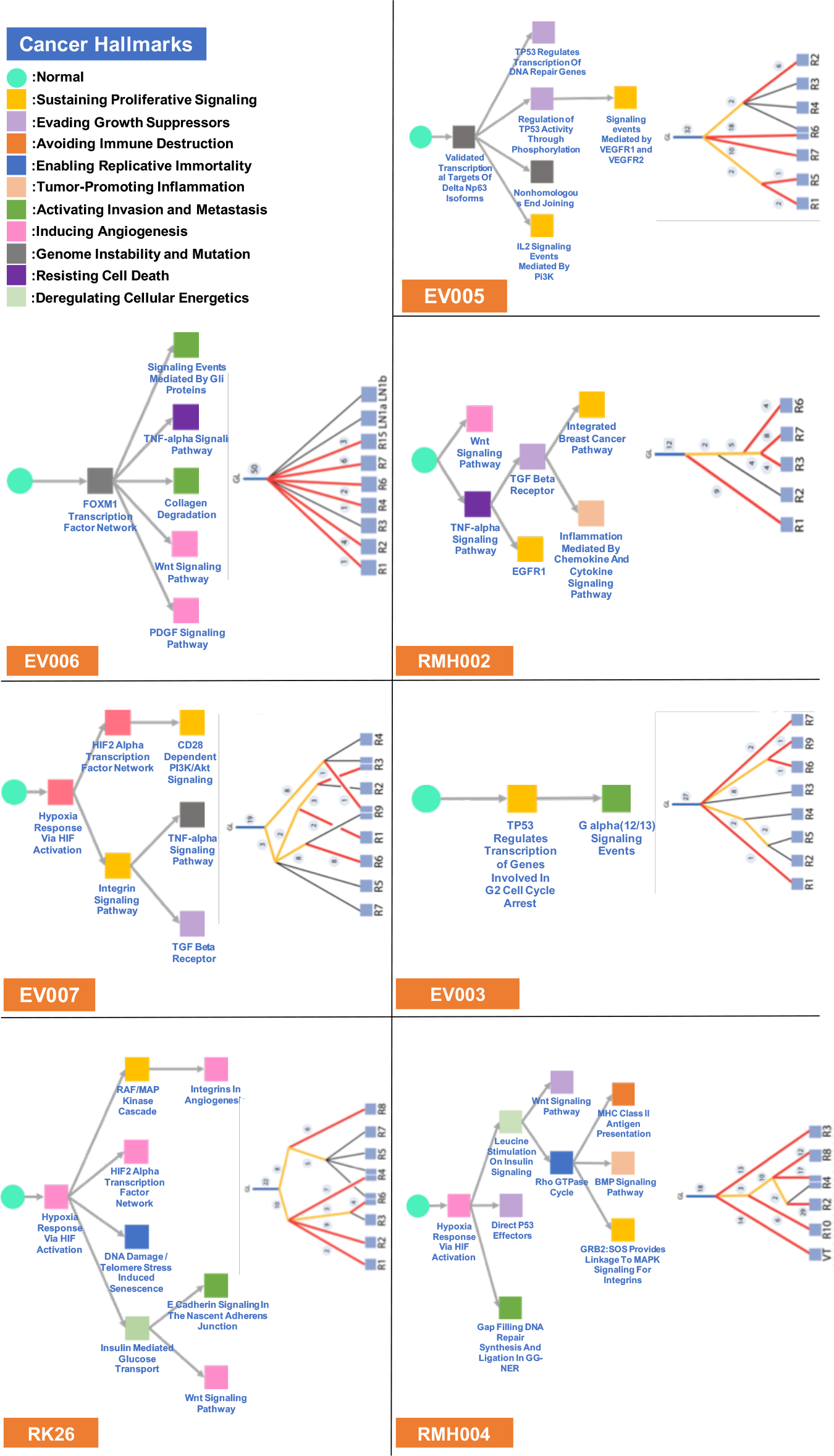
ccRCC progression models of acquisition of cancer hallmarks (left side in the panels) with skeletons from LICHeE (right side in the panels and the figure is the same in [3]). The circle and square shapes indicate the normal cell population and subclone population in the proposed progression model, respectively. Cancer hallmarks are represented as colors shown in the top left panel. Pathway names are described below the box.

**Table 1.** Counts of subclones (patients) with each cancer hallmark. For example, “10 (7)” in Sustaining Proliferative Signaling means that 10 subclones have the cancer hallmark, and it was observed in 7 patients. Columns “trunk” and “private” list the numbers of cancer hallmarks in the common ancestral subclone and in subclones without any descendants, respectively.

| Cancer hallmarks | Total number | Trunk | Private | Other |
| --- | --- | --- | --- | --- |
| Sustaining Proliferative Signaling | 10 (7) | 1 (1) | 5 (5) | 4 (1) |
| Evading Growth Suppressors | 6 (4) | 0 (0) | 2 (2) | 4 (2) |
| Avoiding Immune Destruction | 1 (1) | 0 (0) | 1 (1) | 0 (0) |
| Enabling Replicative Immortality | 2 (2) | 0 (0) | 1 (1) | 1 (1) |
| Tumor-Promoting Inflammation | 2 (2) | 0 (0) | 2 (2) | 0 (0) |
| Activating Invasion and Metastasis | 7 (5) | 0 (0) | 6 (5) | 1 (0) |
| Inducing Angiogenesis | 12 (6) | 4 (4) | 6 (4) | 2 (1) |
| Genome Instability and Mutation | 4 (3) | 2 (2) | 2 (1) | 0 (1) |
| Resisting Cell Death | 4 (4) | 0 (0) | 3 (3) | 1 (1) |
| Deregulating Cellular Energetics | 3 (3) | 0 (0) | 0 (0) | 3 (3) |

Next, we count the cancer hallmarks observed in the common ancestral subclones (trunk) and subclones without any descendants (private) as shown in Table 1.

The patterns of cancer hallmark acquisition still seemed diverse among patients; however, there were several patterns related to phenotypes when we focused on the trunk and private subsets. In the trunk subset, the “Inducing Angiogenesis” (4 subclones were counted) was the most frequently observed cancer hallmark that was due to pathway alterations caused by VHL mutations. The second most frequently observed cancer hallmark was “Genome Instability and Mutation” (2) caused by transcription factor-related aberrations such as the FOXM1 Transcription Factor Network. In the private subset, Sustaining Proliferative Signaling (5) and Activating Invasion and Metastasis (6) were the most common events among the patients (5 out of 8 patients). In particular, untreated patients (RMH004, RMH008, and RK26) showed Activating Invasion and Metastasis (6).

We also determined the frequency of evolutionary paths of cancer hallmarks up to 2 descendants. The most frequent path is “Normal - Inducing Angiogenesis - Deregulating Cellular Energetics” (3) and the second most frequent paths are “Normal - Inducing Angiogenesis - Inducing Angiogenesis” (2), “Normal - Genome Instability and Mutation - Inducing Angiogenesis” (2), “Normal - Genome Instability and Mutation - Activating Invasion and Metastasis” (2), and “Normal - Genome Instability and Mutation - Evading Growth Suppressors” (2). These results give us biological and clinical implications beyond the SSNVs.

## 4 Conclusion

We developed a method for constructing personalized tumor progression models in terms of cancer hallmark acquisition and demonstrated the effectiveness of this model in terms of interpreting cancer evolutionary trees by means of an actual ccRCC dataset. In the example of ccRCC, identification of druggable target subclones that evolved after pathway alteration (HIF activation with a VHL mutation) is a clinically important problem. Our model has some implications. A cancer hallmark can help us to reduce complexity of cancer development and to characterize the phenotypes of subclones. Our method effectively incorporates cancer hallmarks into the current state-of-the-art tree reconstruction method of evaluation of cancer subclonal evolution. As a future challenge, we will build a unified pipeline for constructing skeletons and estimating confident pathway alteration probabilities. We are extending this approach to other cancer types to reveal the phenotypic features of cancer evolution.

